# Inhibition of BMP and FGF signaling prior to wound epithelium formation leads to an aberrant regenerative response in teleost fish *Poecilia latipinna*

**DOI:** 10.1101/2021.01.27.428424

**Authors:** Isha Ranadive, Sonam Patel, Siddharth Pai, Kashmira Khaire, Suresh Balakrishnan

**Author notes:** Corresponding author: Postal address: Prof. Suresh Balakrishnan, Department of Zoology, Faculty of Science, The Maharaja Sayajirao University of Baroda, Vadodara 390002, Gujarat, India.

## Abstract

The BMP and FGF pathways play a pivotal role in the successful regeneration of caudal fin of teleost fish. Individual inhibition of these pathways led to impaired caudal fin regeneration until the pharmacologic inhibitor of FGF (SU5402) and BMP (LDN193189) were metabolized off. Therefore, in the current study both these pathways were inhibited collectively wherein inhibition of BMP and FGF during the wound epithelium formation led to stalling of the process by bringing down the established levels of *shh* and *runx2*. In members of the treatment group, it was observed that, each blastema grows crouched rather than linear and the regrown lepidotrichia therefore remain tilted down. Amongst the other irregularities observed, the transition from epithelial to mesenchymal cells was found hindered due to down-regulation of *snail* and *twist*, brought about by BMP and FGF inhibition. Compromised expression of *Snail* and *twist* deranged the normal levels of cadherins causing disruption in the transition of cells. Lastly, blocking BMP and FGF delayed blastema formation and proliferation due to diminished levels of *fgf2, fgf8, fgf10* and *bmp6*, while *casp3* and *casp9* levels remained heightened causing accelerated cell death. This study not only highlights the axial role of BMP and FGF pathways in regeneration but also accentuates the collaboration amongst the two. This ingenious coordination of signalling further reinforces the involvement of relaying messenger molecules between these crucial pathways.

**Summary Statement:** BMP and FGF collectively control the process of blastema formation in fish and inhibiting them prior to wound epithelium stage leads to irrevocable damage to the regenerating caudal fin.

## Introduction

More than 200 years ago, Broussonet (1786) reported that adult fish could completely regenerate its fins after amputation. A series of studies from then on, revealed that, the teleost fish which can regenerate a number of body parts including scales, heart muscles (Poss et al., 2002; Raya et al., 2004), spinal cord (Becker et al., 2004), retina (Bernhardt et al., 1996), optic nerve (Becker and Becker, 2002), liver (Sadler et al., 2007) and fin (Johnson and Weston; 1995). This makes them an excellent model for regeneration studies. Amongst the various regenerating organs of the teleost fish, the caudal fin has been a congruous choice for procedures such as amputation and imaging since amputation of the caudal fin does not seemingly affect the survival of the fish in any manner and allows easy monitoring of the fin re-growth (Azevedo et al., 2011; Pfefferli and Jaźwińska, 2015). In a region of oriental climate like Indian subcontinent *Poecilia latipinna* has been used extensively as model for studying fin regeneration (Rajaram et al., 2017).

The caudal fin of teleost is a multi-tissue structure wherein lepidotrichia forms the major framework. Each lepidotrichium consists of two hemi-rays which enclose the mesenchymal tissue, nerves as well as fibroblast cells and to allow finer and more articulated movements of the fin, the ends of the lepidotrichia are branched into fine hair like structures known as actinotrichia (Tal et al., 2010; Münch et al., 2013). The entire skeletal frame is highly flexible and consists of inter-ray tissue which controls the movement of the fin. Upon amputation, caudal fin is fully restored by three weeks in *P. latipinna* (Patel et al., 2019a). This is achieved by epimorphic regeneration involving, in the order of occurrence, wound healing, blastema formation and differentiation. Following amputation, wound closure occurs by 12hpa (hours post-amputation). Wound healing is completed by 24hpa and the wound epithelium stratify further to form apical epithelial cap (AEC) by 48hpa (Patel et al., 2019a). Beneath the AEC, a mound of proliferating cells called blastema emerges which can be observed at 60hpa (Murawala et al., 2017). Blastema formation is a climacteric step of epimorphic regeneration (Lee et al., 2009). It occurs either by dedifferentiation of existing unipotent cells to multipotent cells or by the transformation of epithelial cells into mesenchymal like pluripotent cells. Both of these processes eventually form a pool of mononucleated blastema cells (Sousa et al., 2011; Stewart and Stankunas, 2012). The blastema formation and its maintenance are very crucial for the successful realization of regeneration.

Major processes involved in the blastema formation and its maintenance include the remodelling of the extracellular matrix under the influence of MMPs (Santosh et al., 2011) and the transition of cells from epithelial state to mesenchymal state (Sader et al., 2019). Additionally, a high rate of proliferation is necessary to sustain the pool of blastema cells (Poleo et al., 2001). At the epithelial-mesenchymal interface along the proximo-distal axis, scleroblasts differentiate within the blastema and hence the formation of new dermal bone (lepidotrichia) occurs (Quint et al., 2002). Blastemal cells maintain a growth zone underneath the AEC by regulating cell cycle turn over wherein regulation of apoptosis is also crucial (Azevedo et al., 2011). Gene expression studies have indicated that blastema-epidermis interactions are required for dermal bone patterning during zebrafish caudal fin regeneration (Laforest et al., 1998). The existing extracellular matrix (ECM) also undergoes remodelling and plays a dynamic role during blastema formation (Govindan and Iovine, 2015). Such ECM remodeling might facilitate epithelial-mesenchymal interactions between AEC and blastema.

These cellular processes involved in formation and maintenance of blastema are regulated by various molecular signalling pathways. Epithelial-mesenchymal interactions regulate fin regeneration by multiple surrounding factors such as fibroblast growth factor (FGF), Wnt, bone morphogenetic protein (BMP), Insulin like growth factor, retinoic acid and Notch (Lee et al., 2009). Shibata et al. (2016) have revealed that Fgf3 serves as a signal that promotes the regenerative proliferation of blastemal cells during zebrafish caudal fin regeneration. BMP and FGF pathways have been implicated in both tailbud patterning and tail regeneration of *Xenopus laevis* (Beck et al.,2003, 2006). Cooperative inputs of FGF and BMP (Fgf2+Fgf8+Bmp7) signaling have been shown to induce tail regeneration in urodele amphibians (Makanae et al., 2016). However, the details about cross-talk between these pathways in-depth are yet to be discovered. Therefore, both BMP and FGF signaling were targeted with the application of specific inhibitors in the current study to ascertain their collective involvement in blastema formation.

## MATERIALS AND METHODS

### Animal care, handling and maintenance

Sailfin molly (*Poecilia latipinna*) (Lesueur, 1821) of about the size 4.721 ± 0.340 cm and weight about 2.890 ± 0.696 g were purchased from a local supplier (Oscar aquarium, Vadodara, India) and contained in glass aquaria. They were maintained in the temperature range of 28 ± 2°C. Aquarium water was changed on every alternate day with UV-purified water (Rajaram et al., 2016; Murawala et al., 2018; Ranadive et al., 2019). The behavior of the fish was observed on a daily basis. Handling of fish was carried out according to the ethical principles (Drugs and Cosmetics Rule 1945) approved by the Institutional Animal Ethics Committee (IAEC) constituted as per the guidelines of Committee for the Purpose of Control and Supervision of Experiments on Animals (CPCSEA) (Protocol no. MSU-Z/IAEC/14-2017), India.

### Drug administration and tissue collection

Fish were acclimatized for seven days prior to dozing. After acclimatization, they were divided into three groups named ‘normal control’ (NC), ‘vehicle control’ (VC) and ‘treated’. Each group had six animals for individual methodology. Caudal fins of all the animals from all three groups were amputated marking the 0hpa stage. SU5402 was dissolved into 0.01% of DMSO whereas LDN193189 was dissolved in physiological saline (0.6 N). Fish of normal group were allowed to regenerate till 60 hpa. The vehicle control group received intraperitoneal injections of 0.01% DMSO at 12 hpa and normal saline at 24hpa. The test group, based on the previous studies from the lab, was administered with 2.5 mg/kg of body weight SU5402 at 12 hpa (Saradamba, 2012) followed by 2.5 mg/kg of body weight LDN193189 at 24hpa (Rajaram et al., 2016). The injection volume was adjusted to 25 μl.

Drugs were administered through intraperitoneal injection. Tissue collection was done at 60 hpa for various experiments as mentioned in detail by Murawala et al. (2018).

### Morphometry

Morphology of the caudal fin (n=6 for each group) was observed at 60 hpa for all the three groups using a Leica DM2500 light microscope (Leica, Germany). The regenerative outgrowth was measured under the Leica DM2500 microscope fitted with an inbuilt ocular micrometer.

### Skeletal staining

Caudal fins from all three groups were collected at 60 hpa and were fixed in 4% paraformaldehyde for 1 hour at RT (room temperature) followed by incubation in 100% ethanol at 4°C overnight. Following this, the fins were stained with staining solution (0.1% alcian blue prepared in 30% acetic acid and 0.1% alizarin red dissolved in 70% ethanol) for 5 hours in the dark, at RT. The tissues were then transferred to 100% ethanol overnight at 4°C. Next day, these fins were rehydrated in distilled water for 1 hour at RT following which the fins were transferred in 1% KOH for 5-8 min at RT. Following this, the tissues were transferred into 20% glycerol prepared in 0.8% KOH for 20 min at RT followed by incubation into 50% glycerol (prepared in 0.5% KOH) for 20 min at RT. Subsequently, the tissues were transferred to 80% glycerol (in 0.2% KOH) for 20 min at RT. Finally, the tissues were transferred to 100% glycerol (Ferretti and Geraudie 1995; Laforest et al. 1998). Images of the fins were captured using Lieca DM2500 light microscope (Leica, Germany) and Leica image software.

### Gelatin zymography

Fins were collected in lysis buffer for protein isolation. Total protein was isolated (Patel et al., 2019b; Ranadive et al., 2019) and estimated using Bradford method (Bradford, 1976). 40 μg of total protein was loaded on the 7.5% SDS–polyacrylamide gel with gelatine (5 mg/ml) for electrophoresis and subjected to 100 V for 3 hours at 4°C. Afterwards, the gel was washed twice with 2.5% Triton X-100 buffer for 1 hour each in order to remove the SDS from the gel. Following the wash, the gel was rinsed with incubation buffer twice for 15 min each and then incubated in incubation buffer for 18 hours at 37 ° C for renaturation of the proteins. Finally, the gel was stained with 0.25% Coomassie Brilliant Blue R250 for 4 hours followed by distaining (methanol-acetic acid solution) till clear white bands were seen over a blue background. The band intensities were measured by densitometric analysis using Doc-ItLs software (GeNei, Merck, USA).

### Whole mount BrdU staining

The fishes of all three groups – control, vehicle control and treated (n=3 per group) were incubated in the sepeate fish tank containing dissolved BrdU (25 μg/L) (Sigma-Aldrich, USA) for 24 hours before tissue collection. The fin regenerates were collected at 60 hpa, and the zone of interest was fixed in 4% paraformaldehyde followed by blocking the tissue with 10% BSA prepared in PBS for 1 hour at RT. Further, tissues were incubated in primary antibody (0.15μg/ml of Mouse Anti-BrdU in 5% BSA) (Cat. no. B2531; Sigma-Aldrich, USA) overnight in a moist chamber at 4°C in the dark. Next day, tissues were washed with PBS (three times, each for 5 min) and incubated with FITC (Fluorescein isothiocyanate) conjugated secondary antibody (0.5 μg/ml of Goat Anti-Mouse IgG-FITC in 5%BSA) (GeNei, Merck, USA) (Cat.no. AQ3030F) for 90 min at RT. Following this, they were washed again in PBS (three times, each for 5 min) and mounted with antifade solution (Cat.no. S7114, Sigmsa-Aldrich, Merck, USA). Images were acquired using a Leica DM 2500 fluorescent microscope (Leica, Germany).

### qRT-PCR

Total RNA was isolated from the 60 hpa tissues (n=6) from each group using TRIzol reagent (Applied Biosystems, USA). One microgram of total RNA was reverse-transcribed to cDNA using a one-step cDNA Synthesis Kit (Applied Biosystems, USA). Primers used in the study were designed using PrimerBlast tool of NCBI (Table 1). Quantitative real-time PCR was performed on a LightCycler 96 (Roche Diagnostics, Switzerland) with the following program: 100 s at 95°C followed by 45 cycles (each cycle of 10 s at 95°C, 30 s at 60°C and 30 s at 72°C). Gel electrophoresis and melt curve analysis were used to confirm specific product formation. 18S rRNA gene was used as an endogenous control. Fold change values were calculated using the Livak method (2^-ΔΔCq^) given by Livak and Schmittgen, 2001. Fold change values of the treatment group were normalized to the respective fold change values of the control group. qRT-PCR for each gene was performed in triplicate (technical replicates) with a pooled sample from six fishes (biological replicates).

**Table 1:**
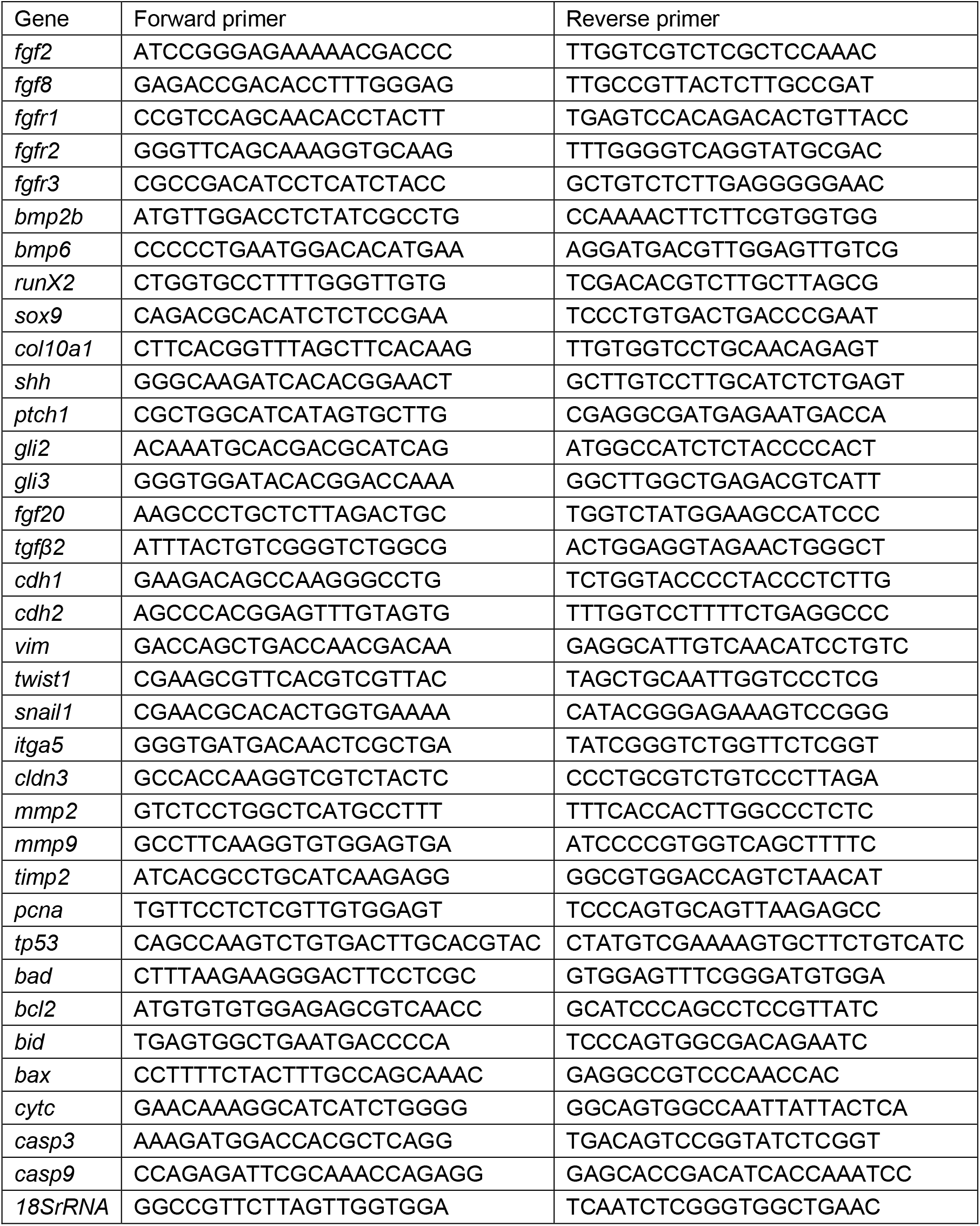
Primer sequences obtained from nucleotide database of NCBI

### Western blot

Regenerated tail fin tissues at 60 hpa were collected, pooled (n=6) and homogenized in Tris-SDS lysis buffer with protease inhibitor (Sigma-Aldrich, USA). 10% homogenates were assayed for total protein content by Bradford method (1976). A total of 25 μg protein was electrophoresed on a 12 % SDS-PAGE gel (Laemmli, 1970). Subsequently, the proteins were electrophoretically transferred on to a nitrocellulose membrane (0.45 μm pore size) at 100 mA for 25 min. The transfer was confirmed by reversible staining using 0.1% ponceau (prepared in 5 % acetic acid). The membrane was probed for β-actin, GRB7, p-Smad1/5, FGF2, Shh, Runx2, E-Cadherin, N-Cadherin, MMP2, MMP9, PCNA, PI3K, AKT, BCL-2 and cleaved Caspase-3 using IgG antibodies (0.1 μg/ml) raised in rabbit, mouse or goat. Biotinylated goat anti-rabbit, mouse anti-goat, and rabbit anti-mouse IgG was used as the secondary antibody (0.5 μg/ml). Streptavidin conjugated ALP-based staining method was used for detecting the protein of interest as a dark band against a white background. β-actin was used as loading control.

### DNA fragmentation

DNA was isolated from fin tissue (n=6 animals per group) as mentioned in detail by Patel et al., 2019. Briefly, 25 mg of tissue was collected in 1 ml of SNET buffer (Sambrook and Russell, 2006) and was incubated at 37°C until the solution turned milky white. Following this, 25:24:1 phenol: chloroform: isoamylalcohol was added to the vial in 1:1 ratio and was incubated on a rocking platform at RT for 30 min. The mixture was then centrifuged at 15000 g for 15 min at 4°C. The aqueous phase was transferred to a new vial and isopropanol was added in equal ratio. DNA precipitation was achieved by centrifuging the vial at 15000 g for 15 min at 4°C. The pellet was washed with 70 % ethanol. Pellet was air dried and dissolved in 30 μl of distilled water. DNA was quantified using UV-VIS spectrophotometer at 260 nm (Shimadzu UV-1800, Japan). 50 ng of DNA from each group was loaded in 1 % agarose gel. Agarose gel electrophoresis was carried out at 100 V for 30 min. The gel image was taken by GeNei imaging system (Merck, USA).

### Statistical analysis

The data of morphometry was expressed as mean ± standard deviation whereas other results were expressed as the mean ± standard error of the mean. Statistical significance was determined either by unpaired Student’s t-test or One-Way ANOVA using GraphPad Prism 5 (San Diego, CA, USA). A *p* value less than or equal to 0.05 was considered statistically significant in all cases.

## Results

### Dual inhibition of BMP and FGF pathways delays and alters blastemal outgrowth

The length of the regenerate was measured for normal control, vehicle control and treatment group (LDN193189+SU5402). The treatment group revealed a remarkable decrease in the length of blastema when compared to both NC and VC (Table 2). To determine the inhibition of BMP as well as FGF signaling western blot for p-Smad1/5 and GRB, downstream molecules of respective pathways, was performed. The results confirmed the shutting down of the pathway which can be seen by diminished levels of p-Smad1/5 and GRB in the treatment group compared to both the control groups (Fig. 1H, Table 3). The images for fins were captured before (Fig. 1A) and after amputation (Fig. 1B). Subsequent to the amputation, images were taken at 12hpa (dosing stage for SU5402) showing the closure of the wound (Fig. 1C). At 24hpa, a thin layer of wound epithelium (dosing stage for LDN193189) can be observed in Fig. 1D. At 60 hpa, the peculiar protrusion of proliferating cells called blastema can be observed for normal control (Fig. 1E) and vehicle control group (Fig.1F) whereas the dual inhibition group exhibited anomalies like bending of lepidotrichia and reduction in length of regenerate (Fig. 1G). Moreover, the blastema in the treated group was formed in a conspicuous downcurved manner covered with a thin epidermis.

**Table 2:** Morphometric analysis of caudal fin regenerate at 60 hpa (n=6). Values are expressed as mean ± SD. Values with same superscript are not statistically significant.

| Group | Length of the regenerate |
| --- | --- |
| Normal control | 11.42 ± 1.12<sup>a</sup> |
| Vehicle control | 10.58 ± 0.98<sup>a</sup> |
| Treated (LDN193189+SU5402) | 4.25 ± 0.39<sup>b</sup> |

**Table 3:** Quantification of western blots by densitometry. Relative band intensities were normalized to the intensity of β-actin of the respective sample. Values are represented in arbitrary units. Values with same superscript are not statistically significant. Superscript “b” and “c” denotes p ≤ 0.05 and p ≤ 0.001 respectively. Values are expressed as mean ± s.e.m

| Protein | Normal control | Vehicle control | LDN193189+SU5402 |
| --- | --- | --- | --- |
| GRB7 | 38.59 ± 2.55<sup>a</sup> | 35.48 ± 2.89<sup>a</sup> | 16.52 ± 0.85<sup>b</sup> |
| p-Smad1/5 | 27.25 ± 1.02<sup>a</sup> | 25.78 ± 1.12<sup>a</sup> | 12.33 ± 0.52<sup>c</sup> |
| FGF2 | 55.13 ± 2.58<sup>a</sup> | 48.19 ± 2.14<sup>a</sup> | 13.58 ± 0.43<sup>c</sup> |
| Shh | 22.36 ± 0.98<sup>a</sup> | 21.96 ± 0.67<sup>a</sup> | 7.52 ± 0.13<sup>c</sup> |
| Runx2 | 32.13 ± 1.06<sup>a</sup> | 33.66 ± 2.35<sup>a</sup> | 11.86 ± 0.59<sup>c</sup> |
| MMP2 | 25.63 ± 1.02<sup>a</sup> | 24.03 ± 0.98<sup>a</sup> | 10.89 ± 0.63<sup>c</sup> |
| MMP9 | 38.23 ± 1.36<sup>a</sup> | 37.11 ± 1.14<sup>a</sup> | 15.87 ± 0.35<sup>c</sup> |
| E-Cadherin | 47.23 ± 2.12<sup>a</sup> | 46.39 ± 1.89<sup>a</sup> | 47.56 ± 2.34<sup>a</sup> |
| N-Cadherin | 27.87 ± 0.88<sup>a</sup> | 26.92 ± 1.03<sup>a</sup> | 11.38 ± 0.56<sup>c</sup> |
| PCNA | 62.38 ± 3.56<sup>a</sup> | 63.57 ± 3.17<sup>a</sup> | 12.62 ± 0.33<sup>c</sup> |
| p53 | 36.15 ± 1.09<sup>a</sup> | 35.43 ± 1.87<sup>a</sup> | 10.92 ± 0.37<sup>c</sup> |
| Cl. Caspase 3 | 48.63 ± 2.47<sup>a</sup> | 47.32 ± 2.04<sup>a</sup> | 66.87 ± 3.77<sup>b</sup> |
| Bcl-2 | 32.96 ± 0.99<sup>a</sup> | 29.58 ± 0.78<sup>a</sup> | 11.47 ± 0.52<sup>c</sup> |

**Fig. 1:**
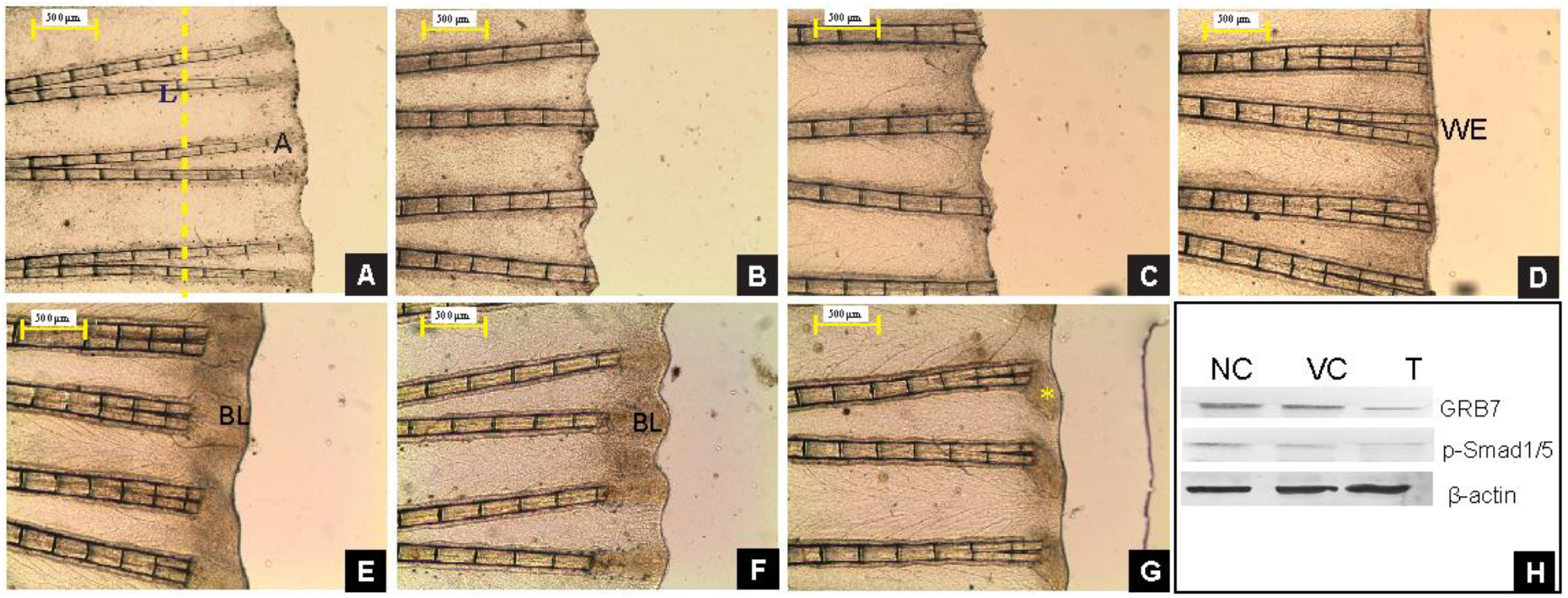
Morphological observation upon inhibition of BMP and FGF pathway. A) Represents the uncut fin where L depicts the lepidotrichium and A is for actinotrichium along with yellow dotted line representing the plane where amputation will be made; B) shows the image of fin on amputation at 0hpa with the yellow dotted line showing the amputation plane; at 12hpa a single layer of epithelium covers the wound which is visible in C), it also represents the stage at which SU5402 was dosed while D) depicts the 24hpa in fin when LDN193189 was dosed intraperitoneally, WE stands for wound epithelium stage; E) and F) are the normal control and vehicle control fins at 60 hpa which marks the blastema (BL) stage; G) dual inhibited fin shows bent blastema marked with an asterisk; H) Western blot of GRB and p-Smad1/5, the downstream molecules of BMP and FGF pathways wherein β-actin is the endogenous control.

### Inhibition of BMP and FGF pathway leads to skeletal deformity in caudal fin

To observe the fin skeleton, alcian blue-alizarin red staining was performed. The results obtained depicts the blue colored area (cells contributing to formation of actinotrichia) in the normal (Fig. 2A) and vehicle control animals at blastema stage (Fig. 2B). The lepidotrichia are stained red in color. The treated group, on the other hand, showed no such signs of cartilage condensation (which is normally stained by alcian blue colour) in the deformed blastema (Fig. 2C). The fishes in the treated group were observed beyond the blastema stage and the results revealed unusual fin skeleton formation. In 9 dpa regenerate, the bent actinotrichia can be seen (Fig. 2D), later these actinotrichia were seen to achieve the normal length but however they failed to ossify. Therefore, the newly formed lepidotrichia at the amputation plane can be seen tilted down even in 14 dpa (Fig. 2E) and 25 dpa (Fig. 2F) fin tissues. Apart from this observation, bifurcation of the regenerate was also delayed wherein the furrow never progressed towards the original fin (Fig. 2D-F). In order to substantiate these observations, genes responsible for patterning of the fin formation were studied for their differential expression. Results of real-time PCR revealed that the expression of *fgf2, fgf8, fgfr1, fgfr2, fgfr3, bmp2b, bmp6, runx2, sox9, col10a1, shh, ptch1, gli2* and *gli3* were completely down-regulated in the LDN193189+SU45402 treatment group (Fig. 2G). To reaffirm the findings of transcript levels, western blot was performed for few key molecules like FGF2, Shh and Runx2 and the experiment revealed decreased levels of these proteins (Fig. 2H, Table 3).

**Fig. 2:**
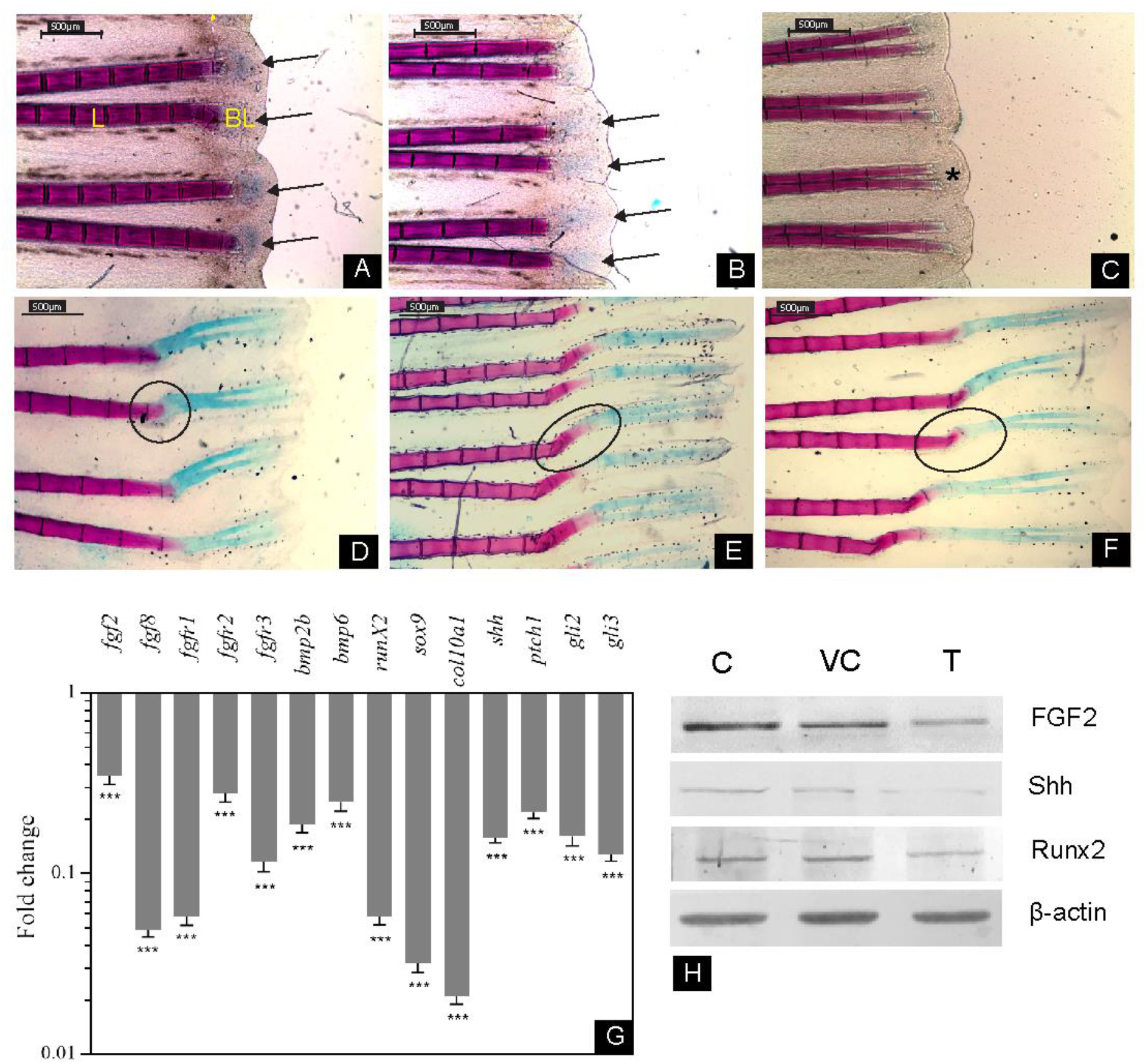
Structural and molecular analysis of fin ray patterning. A-F shows the alcian blue-alizarin red stained fins wherein A) and B) represents the normal control and vehicle control fins at 60 hpa with blue colored region visible in the blastema while C) shows no such coloration in the SU5402 and LDN193189 dosed fins represented by asterisk; D), E) and F) are treated fins allowed to regenerate and images were captured at 9dpa, 14dpa,and25 dpa respectively; G) is the Relative transcript level expression of genes involved in patterning. Fold change values for the treated group was normalized with the values of the control fin. Error bars represent standard error of mean and asterisk depicts p value where * highlights p≤0.05, ** marks p≤0.01 and *** stands for p≤0.001;(n=6); H) depicts the western blot for FGF2, SHH, and Runx2 wherein β-actin was used as the loading control.

### Dysregulated ECM remodeling hampers epithelial-mesenchymal transition upon inhibition of BMP and FGF pathway

In order to initiate as well as continue blastema formation, the tissue underneath the newly formed wound epithelium has to undergo ECM remodeling to provide space for proliferating cells. Thus, specific key regulators involved in ECM remodeling were studied at their transcript levels. Expression of *cdh1* was found to be upregulated in treatment group whereas *cdh2* and *vim* levels were decreased (Fig. 3A). Concurrent results were obtained for E-Cadherin and N-Cadherin at the protein level by western blot as well (Fig. 3B, Table 3). Other than these ECM remodeling mediators, MMPs also play a pivotal role in cell migration while blastema formation and hence MMP2 and MMP9 were checked at transcript and protein level. Both *mmp2* and *mmp9* gene exhibited significant down-regulation upon dual inhibition of BMP and FGF signaling pathways (Fig. 3A). The protein expression of MMP2 and MMP9 was also reduced notably for the treated group when compared to control and vehicle control group (Fig. 3B, Table 3). Additionally, activity levels of both the forms (pro- and active) of MMP2 and MMP9 were studied by gelatin zymography. The results indicated decreased activity of both pro and active form of both the gelatinases upon drug administration in comparison to the control and vehicle control group (Fig. 3C, Table 4). Furthermore, TIMP- the well-known inhibitor of MMP, was also studied for its gene expression levels. *timp2* levels were observed to be increased upon dual inhibition (Fig. 3A). To further validate the disruption of EMT, *twist1, snail1, itga5, cldn3, fgf20* and *tgfβ2* were studied for their gene expression levels. The results revealed down-regulation of *fgf20, tgfβ2, twist1, snail 1, itga5* and increased expression of *cldn3* (Figure 3A).

**Fig. 3:**
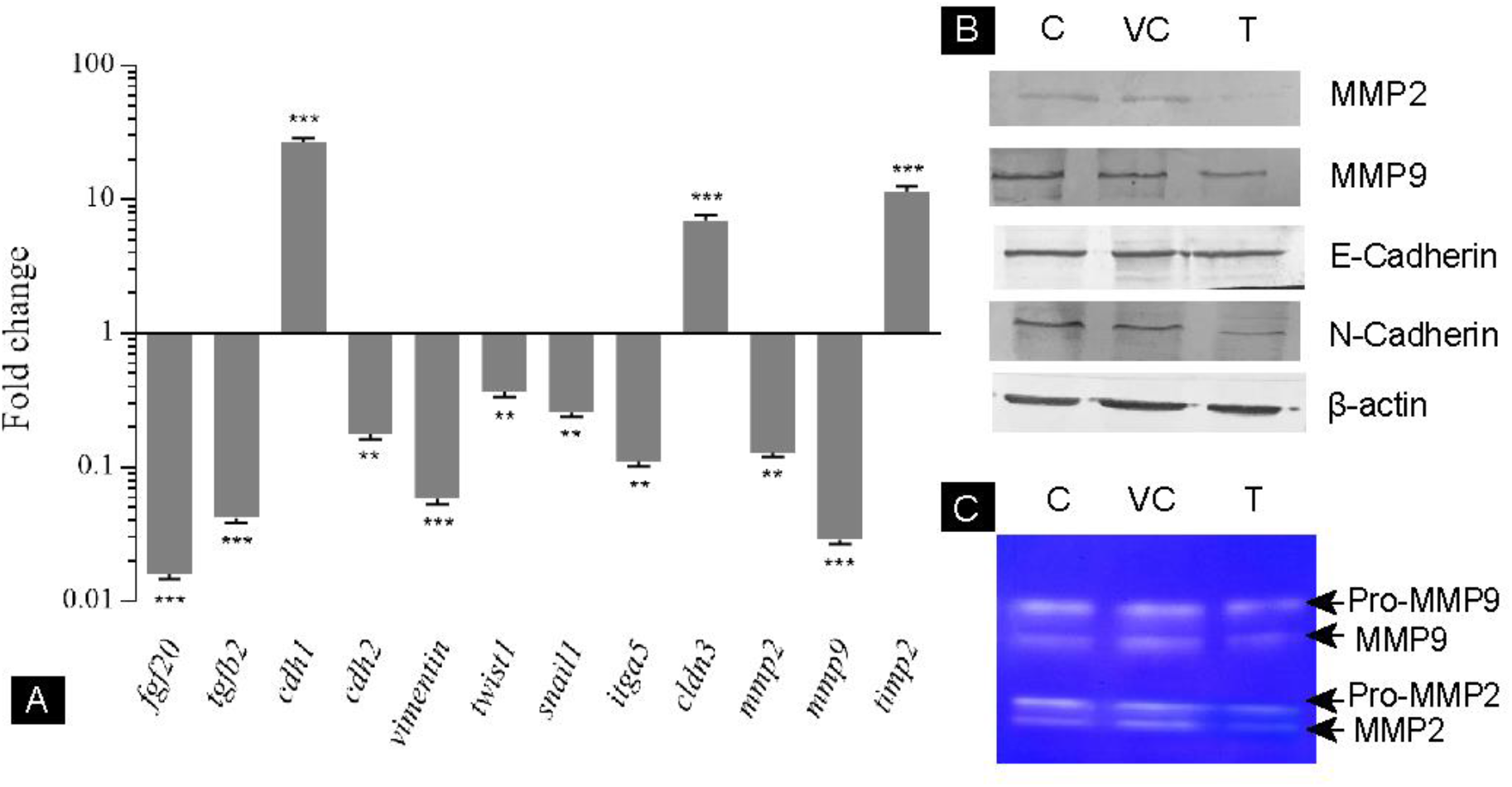
Effect of inhibition of BMP and FGF pathway on EMT. A) Shows the relative transcript level expression of the genes involved in the process of EMT. Fold change values for the treatment group was normalized with the values of the control fin. Error bars represent standard error of mean and asterisk depicts p value where * highlights p≤0.05, ** marks p≤0.01 and *** stands for p≤0.001;(n=6); B) is a representative image of western blot and C) shows the gelatin zymography for the enzymes MMP2 and MMP9 in control and LDN193189 treated group.

**Table 4:**
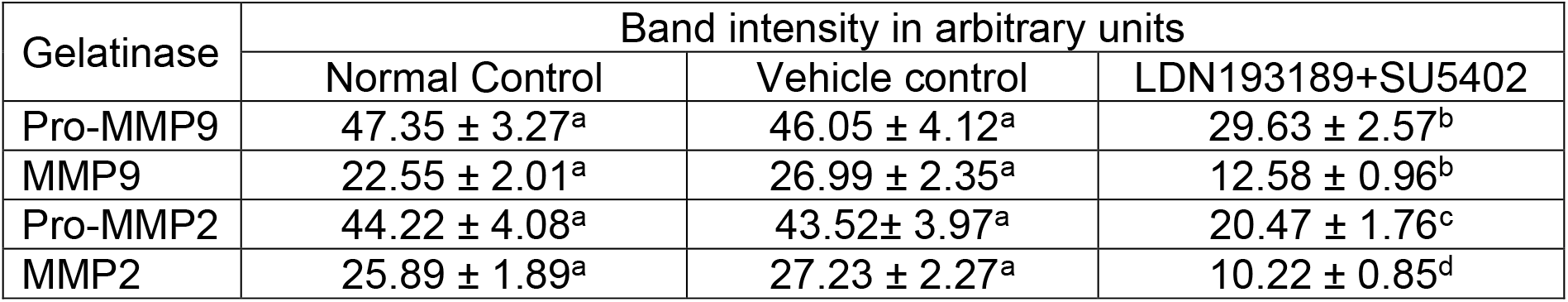
Relative activities of gelatinases represented as band intensities in control and LDN193189+SU5402 treated group fins at 60 hpa (n=6). Values with same superscript are not statistically significant. Superscript “b”, “c” and “d” denotes p ≤ 0.05, p ≤ 0.01and p ≤ 0.001 respectively. Values are expressed as mean ± s.e.m

### Inhibition of BMP and FGF pathways deregulates cell cycle turn over

Cell proliferation at blastema stage was checked by *in vivo* BrdU incorporation assay in the whole fin. Upon comparison of tail fin tissues, it revealed that the proliferating cells are found more profusely in the normal control (Fig. 4A) and vehicle control (Fig. 4B) group than in the treated group (Fig. 4C). Very few proliferating cells can be observed in the AEC and the underlying deformed blastema of treated fin tissues. Also, a weak proliferation rate in the inter-ray tissue of treated animals was recorded compared to controls wherein the proliferating cells were observed in a great abundance (Fig. 4C). The corresponding bright field images were also captured for normal control (Fig. 4D), vehicle control (Fig. 4E) and treated (Fig. 4F) group fin tissues for authenticating the field of vision. After visualizing the reduction in the number of proliferating cells due to dual inhibition, PCNA, a marker of cell proliferation was assessed at mRNA and protein level at blastema stage in both controls and treated animal fin tissues. PCNA was found to be decreased remarkably in the treated group at both gene (Fig. 4G) and protein level (Fig. 4H, Table 3) reaffirming the BrdU assay results.

**Fig. 4:**
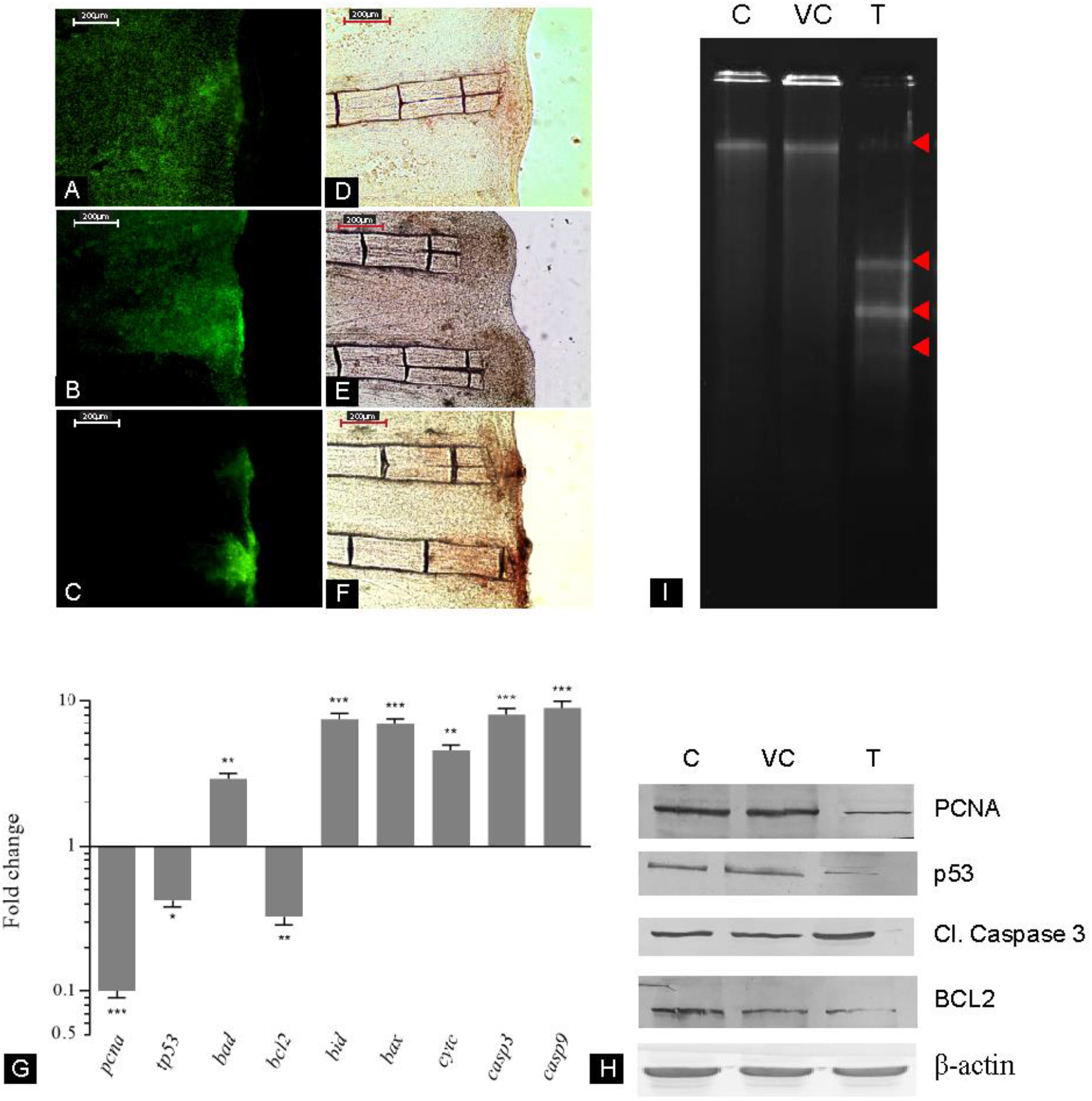
Effect of inhibition of BMP and FGF pathway on cell cycle turnover. A), B) and C) represent the BrdU incorporation in control, vehicle control and LDN193189 treated fins at 60 hpa, while D), E) and F) are the bright field images for the same; G) illustrates the real time PCR for genes involved in proliferation and apoptosis of cell wherein the bars represent the fold change values for the treatment group when compared the to the control fin. Error bars represent standard error of mean and asterisk depicts p value where * highlights p≤0.05, ** marks p≤0.01 and *** stands for p≤0.001;(n=6); H) is the western blot image for PCNA, p53 Cl. Caspase 3 and BCL2 protein wherein β-actin was added as a loading control.

For cell cycle turnover, apoptosis is equally important as cell proliferation. Hence, few markers of apoptosis were also studied to understand the mechanism of cell death induced by inhibition of BMP and FGF pathways. Gene expression levels of *tp53, bad, casp3* and *casp9* were found to be elevated in LDN193189 and SU5402 treated fins, contrary to *bcl2*, an anti-apoptotic gene whose expression was found to be decreased (Fig.4G). Protein expression levels of BCL-2 was found to be decreased too and cleaved Caspase-3 revealed an upregulation following the treatment of LDN193189 and SU5402 (Fig.4H, Table 3). Further, DNA damage being one of the hallmarks of apoptosis was studied via DNA fragmentation assay. LDN193189 and SU5402 treatment resulted in an elevated DNA fragmentation when compared to the control and vehicle control (Fig.4I).

## DISCUSSION

Being an important stage of epimorphic regeneration, the study was aimed to investigate the involvement of BMP and FGF during blastema formation. Upon inhibition of both the pathways, diminished and bent blastema was observed. One of the unique trends observed in this study is the bending of the entire regenerating structure on the disruption of BMP and FGF signaling. This observation was also made in a single inhibition group of LDN193189 treated fishes (Patel et al., 2019b), however in single inhibition group the bending resolves by 14 dpa in dual inhibition group we found that the bending persists even in the later stages of regeneration and hence it indicates that disruption of both the signals is causing this abnormality. This bending is detected from the very beginning viz. early onset of the blastema. Hence it would not be very farfetched to assume that the factors affecting tissue patterning in the regenerate are responsible for the conspicuous bending of the regenerate in the treated group.

Shh is involved in patterning of many structures during the process of regeneration, along with its receptor patched1 (ptch1) and co-factor bone morphogenetic 2b (bmp2b) (Yang et al., 2015; Armstrong et al., 2017). The expression of these genes was found near the stump distal to each lepidotrichium following amputation in zebrafish caudal fin. Shh is expressed in the basal layer of the epidermis at 96 hpa (Armstrong et al., 2017) while ptch1 and bmp2b are located in the matrix secreting scleroblast within the mesenchyme (zebrafish) (Laforest et al., 1998). Once the blastema forms and the cells begin to differentiate, actinotrichium is formed which later constructs lepidotrichium. As the lepidotrichium is shaped, *shh* and *ptch1* expression triggers the bifurcation of the lepidotrichium except for the outermost ones (Quint et al., 2001). The re-expression of *shh* and *bmp2b* in the region of cell proliferation and fresh ossification, suggest their role in the formation and patterning of new dermal bone and potential cell proliferation during fin regeneration (Quint et al., 2001). Our results also exhibited down-regulation of *shh, bmp2b, ptch1* as well as *gli2* at 60 hpa upon inhibition of BMP and FGF pathways which might have led to the abnormal bone formation. As reported by Bai and coworkers (2002), *gli2* induces the expression of *shh*, alteration in which can in turn aberrate the patterning of lepidotrichium. This supports and explains our results, wherein, under the manifestation of LDN193189, BMP signals got derailed and caused disturbed sculpting of lepidotrichium. It is a well-established fact that *gli2* and *gli3*, collectively formulate the skeleton (Mo et al., 1997). Present study reinforces this fact as the exposure of fin regenerate to LDN193189 and SU5402 caused skeletal malformations as expression of gli2 and gli3 were found to be lowered coherently at 60 hpa.

Apart from Gli2 and Gli3 there are reports which suggest that Shh along with FGF2 is required for chondrogenesis and skeletogenesis (Abzhanov and Tabin 2004), the two critical processes essential for actinotrichia and lepidotrichia formation. In our study since FGF pathway was also blocked along with BMP, lowered levels of *fgf2, fgf8* and all the receptors *fgfr1, fgfr2* and *fgfr3* were observed. FGF2 also regulates the expression of Shh as seen in the muscle formation in zebrafish embryo (Yin et al., 2018), suggesting a misregulation of shh in treated fin which persists for a longer time as compared to the single inhibition as reported by Patel et al., 2019b. A study on FGFR2 mutant mice showed bending in the tibial bone along with other abnormal structures which led to Dysplasia (Merrill et al., 2012). Reiterating the same idea, alcian blue and alizarin red staining of the treated fish fin, demonstrated abnormal bending of lepidotrichia, which might have occurred due to the aberrated/inhibited FGF pathway. The observed bone defects in later stages led us to examine the known downstream targets of BMP and FGF signaling during blastema formation, which comprises of bone and cartilage precursor cells. In the fin blastema the major osteogenic transcription factors Runx2a and Runx2b are under the control of BMP signaling (Lee et al., 2000). Both transcript and protein levels of Runx2 were lowered in the treated fins which indicates the inhibition of BMP pathway and also leads to the conclusion that as there is a disruption of signaling observed at 60 hpa and its effects are visible at even later stages as seen in the alcian blue-alizarin red stained fins. Moreover, since fin rays are described as dermal bones, it was observed that the transcription factor associated with cartilage formation, sox9, was down-regulated in the treated fins at 60hpa. Smith and coworkers (2006) have explained the involvement of sox9 and bmp6 in governing chondrogenesis. Current study fortifies this idea as when the bmp levels were found to be lowered at 60 hpa, abnormal cartilage formation was observed in the actinotrichia. Proper activation of FGFR2 and TGF-β is necessary for the recruitment of key tissue patterning regulators (Partridge et al., 2000; Ardi et al., 2009). Corresponding to this concept, inhibition of BMP and FGF pathways, would lead to derailed patterning in the regenerated tissue.

Following structural abnormalities, process of EMT was focused due to its importance in blastema formation. For EMT to occur successfully, digestion and remodeling of the existing extracellular matrix is a must and many factors are responsible for this process (Brown and Badylak, 2014). It is imperative for the cells to dislodge from the basal membrane and migrate onto the amputation plane in order to close the wound surface. To assess the effect of BMP and FGF inhibition on ECM remodeling, gene and protein expression pattern of major factors such as E-Cadherin, N-Cadherin, MMP2, MMP9, TIMP-2, TIMP-3 and TIMP-4 was checked. MMP2 and MMP9 are required for digestion of matrix, but their levels were lowered when BMP and FGF pathways were shut down. FGF2 and FGFR2 are known to induce the proMMP9 to form MMP9 which is the active enzyme observed in mouse angiogenesis model (Partridge et al., 2000; Ardi et al., 2009). Hence, lowered levels of these FGF family members in the caudal fin treated with the compounds might have led to inhibition of MMP9 activity which was confirmed by the gelatin zymography. Both FGF and BMP signaling play a crucial role in lowering of MMP activity. MMP9 has been shown to restrain corneal epithelial migration through the activation of TGFβ3 (Chen and ten, 2016). Buch et al. (2018) have reported that FGF2, FGF8 and FGF10 are required for triggering the MMP2 and MMP9 activity. Hence upon inhibition of the FGF pathway in the current study, it was observed that MMPs were down-regulated and thus it led to improper blastema formation. Once it was established that ECM remodeling was hampered during BMP and FGF inhibition, the possibility of epithelial to mesenchymal transition was lowered. The effect of inhibition of these pathways on EMT was checked by assessing transcript and protein levels of E-Cadherin and N-Cadherin. N-Cadherin which is required for the cells to undergo EMT was significantly reduced along with vimentin. Apart from the markers of EMT, Fgf20 was found to be expressed during initiation of fin regeneration at the epithelial-mesenchymal boundary by Whitehead et al. (2005). Also, FGF2 directly cannot cause down-regulation of N-Cadherin since it recruits FGFs during EMT (Suyama et al., 2002). In fact, it is BMP2 which upregulates N-Cadherin via non-canonical SMAD2/3 pathway (Zhao et al., 2018). These reports affirm the coherent roles of N-Cadherin, BMP2 and FGF2 in transition of epithelial cells to mesenchymal lineage. The diminished levels of N-Cadherin under lack of BMP2 and the resulting reduction of FGF2 levels are thus responsible for the disoriented blastema formation. Furthermore, hiked levels of both, E-Cadherin and cldn3, confirms the prevalent intercellular attachment, additionally preventing EMT. Claudin 3 and Claudin 4 have been reported to promote mesenchymal to epithelial cell transition during ovarian cancer (Lin et al., 2013) and its increase in our study further substantiates the inhibition of EMT.

To reaffirm the inhibition of EMT, SNAIL and TWIST which are the master regulators (Odero-Marah et al., 2018), were studied. The results indicate their lowered levels in the treated group which proves that upon inhibition of BMP and FGF pathway EMT was lowered and hence at 60 hpa a proper blastema was not formed. Itga5 (Integrin α5β1), a fibronectin receptor, is implicated in EMT and wound healing and is more strongly expressed in highly invasive colon cancer cell lines than in poorly invasive ones (Nam et al., 2012). Results obtained in present study demonstrate down-regulation of *itga5* as a function of altered *snail* and *twist* levels, eventually curbing the EMT.

Once the blastema has been formed, its maintenance by continuous proliferation with a regulated amount of apoptosis is necessary. An abnormal shift in the balance in favor of any of these two processes could have lethal consequences, for instance, a cancerous condition in case of the former (Ventura et al., 2007) and atrophic outcomes as an effect of the latter (Dupony-Versteegden, 2005). Nevertheless, in case of a regenerating system, in order to recuperate the loss of the amputated appendage, regrowth is supported by an extremely high rate of cell proliferation in the pool of blastema cells (Patel et al., 2019b), subsequently, as the elongation of the regenerate occurs, blastema cells are utilized for re-differentiation and reorganization into adult tissue (Rajaram et al., 2017). Hence, cell proliferation is important for the formation as well as for the perpetuation of the blastema. Herein, cell proliferation was assessed at both gene and protein level using PCNA as marker. The best understood function of PCNA, till date, is its role in DNA replication (Kelman, 1997). Genetic and protein expression levels of PCNA show marked down-regulation in the treated group as compared to the control one indicating impairment of cell proliferation upon dual inhibition of the FGF and BMP signaling pathways. Down-regulation of PCNA in the treated group was further confirmed with BrdU incorporation assay. Thummel et al. (2006) have reported similar results wherein knockdown of fgfr1 during zebrafish fin regeneration decreased the number of BrdU positive cells of both blastema and regenerative outgrowth.

In order to observe the effect of the dual inhibition on apoptosis, anti-apoptotic marker (*bcl-2*) and pro-apoptotic markers (*tp53, bad, casp3* and *casp9*) were screened for their gene and protein expression levels in control as well as treated group. Down-regulation of the anti-apoptotic gene Bcl-2 in the treated group correlates with the simultaneous upregulation of pro-apoptotic genes *tp53, bad, casp-3* and *casp-9* indicating a higher rate of apoptosis leading to a negative impact on cell cycle turnover. Pellettieri et al. (2010) have also reported the presence of BCL-2 during planarian regeneration. Increased levels of *casp-3* have been reported during blastema formation upon inhibition of BMP pathway using LDN193189 (Patel et al., 2019b). Furthermore, apoptosis was characterized by DNA fragmentation. DNA fragmentation assay was performed in which the treated group showed many fragments of DNA indicating higher apoptosis as compared to control fins where a single band was observed. It has been shown that Caspase-3 is required for DNA fragmentation and other morphological changes of the cell undergoing apoptosis (Jänicke et al., 1998). Our results showed that the inhibition of BMP and FGF pathways paved way for Caspases to get activated and hence apoptosis to occur leading to impairment of blastema and deviant regenerative response.

## Conclusion

The current study was aimed at revealing the interplay between BMP and FGF pathway in the process of blastema formation, a crucial step required towards successful regeneration. Inhibiting BMP and FGF pathways through LDN193189 and SU5402 respectively prior to blastema formation, depicted a small and bent blastema which later on differentiated to form the fin with abnormally bent lepidotrichia. The probable reason for bent lepidotrichia in regenerated fin could be due to lowered levels of *shh* and *runx2*, caused by inhibition of BMP pathway that normally regulates the expression of these molecules along with FGF which recruits *shh* during tissue patterning. Apart from bending of the fin rays, EMT was completely hampered as substantiated by reduced N-Cadherin, MMP2, MMP9, *snail, twist* and *itga5* levels. FGF inhibits the E-Cadherin levels during EMT, however its inhibition led to continued expression of E-Cadherin ultimately restricting the process altogether. Lastly, significantly reduced blastema was observed upon inhibition of BMP and FGF with heightened expression of *casp3* and *casp9* exhibiting increased apoptosis in treated group animals. Surprisingly tp53 was lowered in the treated group and the reason remains to be unknown. Overall, BMP and FGF not only play an important role in the processes like EMT, proliferation, and apoptosis during blastema formation but are also important for inducing proper ray formation in the later stages of regeneration. This has been summarized pictorially in Fig. 5.

**Fig. 5:**
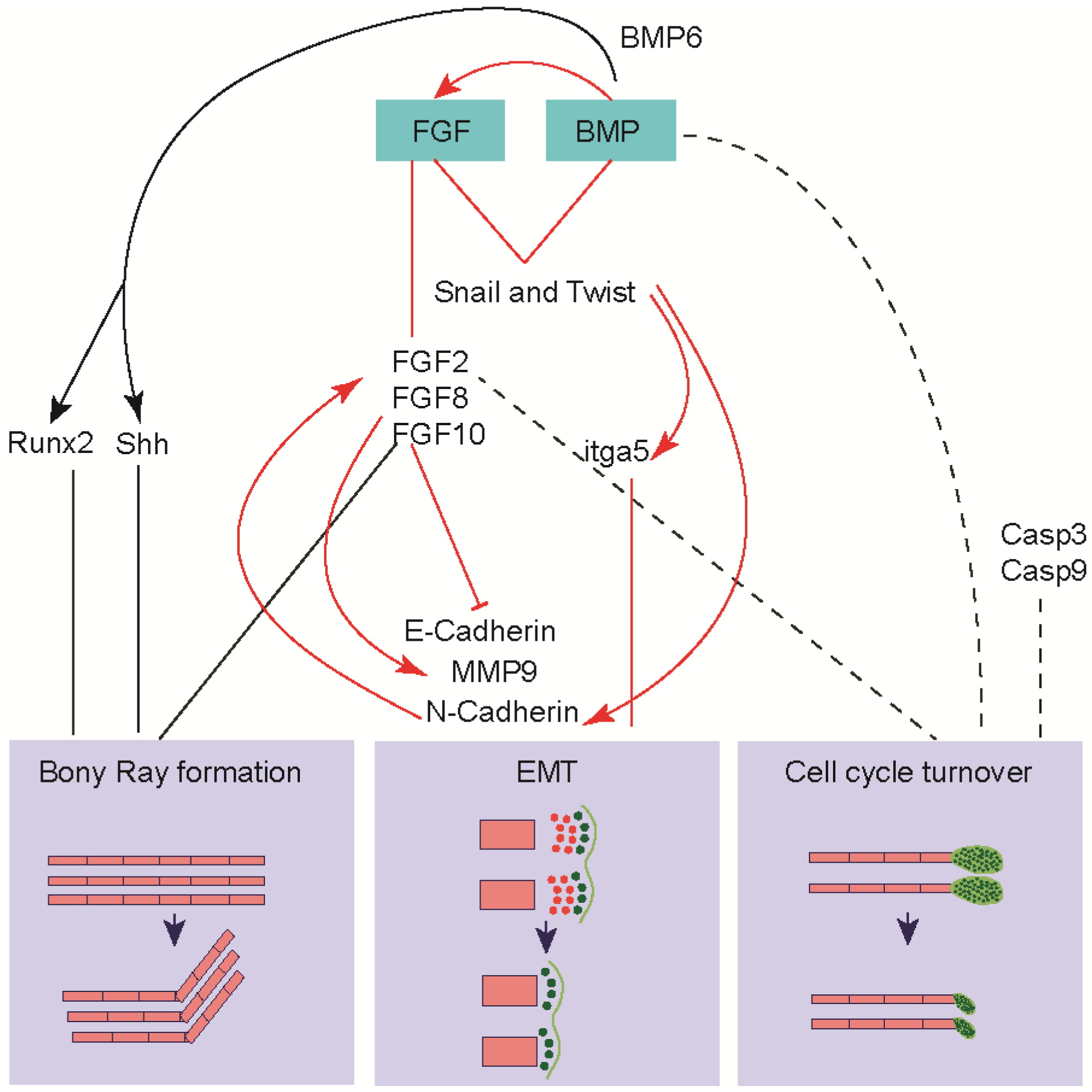
The cross-talk between BMP and FGF pathway during caudal fin regeneration

## ACKNOWLEDGEMENTS

IR and SP are indebted to Science and Engineering Research Board, New Delhi and University of Grant Commission, respectively for support in terms of fellowship.

